# Trial Averaging for Deep EEG Classification

**DOI:** 10.1101/2023.02.09.527905

**Authors:** Jacob M. Williams, Ashok Samal, Matthew R. Johnson

## Abstract

Many signals, particularly of biological origin, suffer from a signal-to-noise ratio sufficiently low that it can be difficult to classify individual examples reliably, even with relatively sophisticated machine-learning techniques such as deep learning. In some cases, the noise can be high enough that it is even difficult to achieve convergence during training. We considered this problem for one data type that often suffers from such difficulties, namely electroencephalography (EEG) data from cognitive neuroscience studies in humans. One solution to increase signal-to-noise is, of course, to perform averaging among trials, which has been employed before in other studies of human neuroscience but not, to our knowledge, investigated rigorously, particularly not in deep learning. Here, we parametrically studied the effects of different amounts of trial averaging during training and/or testing in a human EEG dataset, and compared the results to that of a related algorithm, Mixup. Broadly, we found that even a small amount of averaging could significantly improve classification, particularly when both training and testing data were subjected to averaging. Simple averaging clearly outperformed Mixup, although the benefits of averaging differed across classification categories. Overall, our results confirm the value of averaging during training and testing when single-trial classification is not strictly necessary for the application in question.

**Highlights:**

- Averaging trials can dramatically improve performance in classification of EEG data
- The benefits can be seen when averaging on both training and test datasets
- Simple trial averaging outperformed a popular related algorithm, Mixup
- However, effects of averaging differed across different stimulus categories

## 1. Introduction

Some classification tasks have enough information in the training set to achieve above-chance classification in the test set, but the signal-to-noise ratio is low enough that convergence of the classifier is unreliable given reasonably available time and computational power. For example, in certain datasets, choosing a different train/test split can vastly alter the performance. Additionally, it is frequently observed that an artificial neural network will show no progress toward convergence (e.g., lowered validation cross-entropy) for dozens of epochs at the beginning of training, but may eventually progress toward convergence. This suggests that it can take many permutations of minibatches before the algorithm lands on one that allows continued progress toward improved classification performance. This scenario can be difficult to rectify with most training schedules, whether using early stopping or not.

In some of these tasks, training on same-class averaged signals will allow for more consistent convergence of the classifier. For example, in neuroscience research, working with pre-averaged signals is a somewhat common practice in multivariate pattern analysis (MVPA). Neuroscience data is typically strongly temporally and spatially aligned, at least within a single subject, but often has poor signal-to-noise ratio with regard to the semantic features of interest. While there will always be some trial-to-trial variability due to subject-related factors such as attention, the signals are generally similar enough that averaging can be a viable method to improve the signal-to-noise ratio of the desired semantic features. Thus, training and classifying with a learning algorithm can often be more reliable when working with averaged signals as opposed to single-trial signals.

This training technique may be particularly valuable in circumstances where the amount of data collected is necessarily limited, which is often the case in neuroscience and other biomedical domains. Neuroscience data also have the complication that the underlying ground truth about the quality and theoretical classifiability of a single-trial signal is fundamentally unknown, even to a human observer, unlike other domains such as photographic images or natural language processing. For example, a subject in an electroencephalography (EEG) experiment may be distracted or disobey instructions when they are supposed to be attending to visual stimuli, leading to a signal of extremely poor quality or even of the incorrect underlying semantic class, but indistinguishable from high-quality data in terms of any macroscopic artifacts. Thus, the signal may be unclassifiable through no fault of the algorithm applied to it, but this can be difficult to determine either during or after data collection. This presents an opportunity for improvement using an averaging approach, which can balance out variation in signal quality between trials if occurrences of very poor data quality trials are sufficiently rare.

Averaging also provides a form of data augmentation, with this use of averaging being well-established in the machine learning field. The Mixup algorithm [1] is one well-known and straightforward application of averaging to augment a dataset. Mixup produces virtual feature/label vectors of the form:

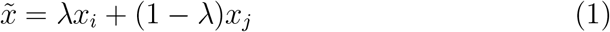

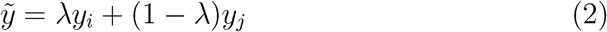

where (*x*_*i*_, *y*_*i*_) and (*x*_*j*_, *y*_*j*_) are feature/label pairs from the dataset and *λ* ∈ [0, 1], sampled from a beta distribution. In essence, Mixup produces weighted averages of training samples and their labels as the augmented data to train on. Using Mixup has been shown to increase network accuracy, particularly in larger, slower-to-converge networks, and to improve robustness to adversarial examples. Notably, the samples chosen for Mixup do not need to be in the same underlying class; the resulting data is a superposition of the two input samples and their labels.

A similar technique, called Sample Pairing, does simple averaging on the pixel intensities of two images from the training set, while only using the label of the first of the two images. While this prevents the training accuracy from being high, it allows for a successive fine tuning without Sample Pairing that results in a reduction of error rate in tasks such as ILSVRC and CIFAR-10 [2]. Other methods have used interpolation and extrapolation in feature space to produce augmented datasets [3, 4]. These methods share with Mixup the qualities of being domain-agnostic and being generated from a combination of existing samples.

Signal averaging before classification has a long history in MVPA. Early MVPA work in functional magnetic resonance imaging (fMRI) used correlational analysis after averaging several runs’ worth of trials [5]. Even more recent work has discussed the use of “supertrials,” or randomly averaged sets of individual trials, for use in EEG pattern analysis [6]. These “supertrials” can be used in either or both of the training and testing phases depending on the structure of the particular study. MVPA tutorials have sometimes discussed the common applications of trial averaging, and the impact of the increased signal-to-noise ratio (see, for example, [7]). However, it is also pointed out that being able to use individual trial information can allow for a more sensitive classifier [8]. To our knowledge, though, to date there has been no formal exploration of the space of averaging, over either or both of the training and testing phases, particularly with regard to deep-learningbased MVPA. Thus, in the present study we focus our efforts on providing such an exploration.

## 2. Material and Methods

### 2.1. Problem Definition

The goal of this research was to examine the effects of averaging trials in a particularly low signal-to-noise electroencephalography dataset. In particular, we sought to address the following research questions:

- How does averaging samples within-class during training affect the performance at test time?
- How does averaging test samples affect the ability to classify the test set?
- Is simple, within-class averaging more or less effective than Mixup?
- Do the effects of averaging differ by stimulus class in this dataset?

### 2.2. Datasets

The EEG dataset we used for this experiment was previously collected by one of the authors in a cognitive neuroscience study [9] and is one that we have also used previously [10] to develop deep-learning methodology for classifying neuroscience data. This dataset was recorded on a 32-channel EEG cap, with a gain of 10,000, a bandpass filter of 0.01-100Hz, stored at 14-bit precision, at a 250Hz sampling rate.

There are two main sets of signals available in this dataset: initial presentation and refreshing within short-term memory. The initial presentation refers to the period of time wherein the subject was viewing a visual stimulus belonging to one of three classes (faces, scenes, or words). The refresh condition refers to a period wherein the subject was prompted to think back to or imagine one of those recently perceived visual stimuli that was no longer visible onscreen. We have used this dataset in developing the Paired Trial Classification technique [10], where the initial presentation period of the task was used to classify the category of visual stimuli as they were perceived. In prior analyses by members of our research group, we had found that the refresh condition was capable of being classified above chance when working with single trials, but not by a very wide margin, and not reliably for all types of analyses [9]. Because that condition pushed the boundaries of what is possible to classify from single-trial human neuroscience data — a brief thought measured with scalp electrodes and activity on the order of microvolts, a very weak signal by the standards of most cognitive neuroscience studies — we considered it an ideal test case for examining how different averagingbased techniques might allow a barely-detectable signal to be classified more reliably.

The refresh dataset consists of data recorded from 37 human participants with approximately 100 trials each, split evenly between the three stimulus categories (classes) before pre-processing and artifact rejection. Each trial comprises 2.1 seconds of data, with the first 100ms forming a pre-trial baseline period, the next 1500ms being the critical period for analysis in which the cue to think back to (visualize) the stimulus was presented, and the final 500ms being a post-cue period.

### 2.3. Neural Network Architecture

We employed a Bidirectional Long Short Term Memory (Bi-LSTM) based architecture due to several recent successes with that architecture in EEG classification, e.g., [11, 12, 13]. As with standard LSTMS, Bi-LSTMs inherently take into account temporal correlations, but are better able to handle long-term temporal correlations by processing the signal from the beginning and the end simultaneously [14]. The depth and number of parameters in the present study’s network were greatly reduced from what is seen in the image classification literature due to the limited availability of data and the low signal-to-noise ratio. Further, an abundance of regularization was necessary to prevent overfitting. The full sequences produced from the Bi-LSTM processing were used as the features to pass to the next layer, rather than simply the final produced output, since the goal was classification rather than prediction of the next time step. Standard hyperbolic tangent activations were used in LSTM layers. Batch normalization was applied to the sequences produced by the LSTM layers to prevent covariate shift and increase generalizability [15]. The LSTM layers were followed by a 1-dimensional average pooling layer to reduce the dimensionality of the produced sequences. Flattening was then applied and the result was fed into dense layers. The dense layers had Leaky Rectified Linear Unit (Leaky ReLU) [16] activations and were followed by dropout and batch normalization layers. The final layer had 3 neurons (one per class) and a softmax activation to produce the ultimate classification decision.

See Figure 1 for the general architecture of the network.

**Figure 1:**
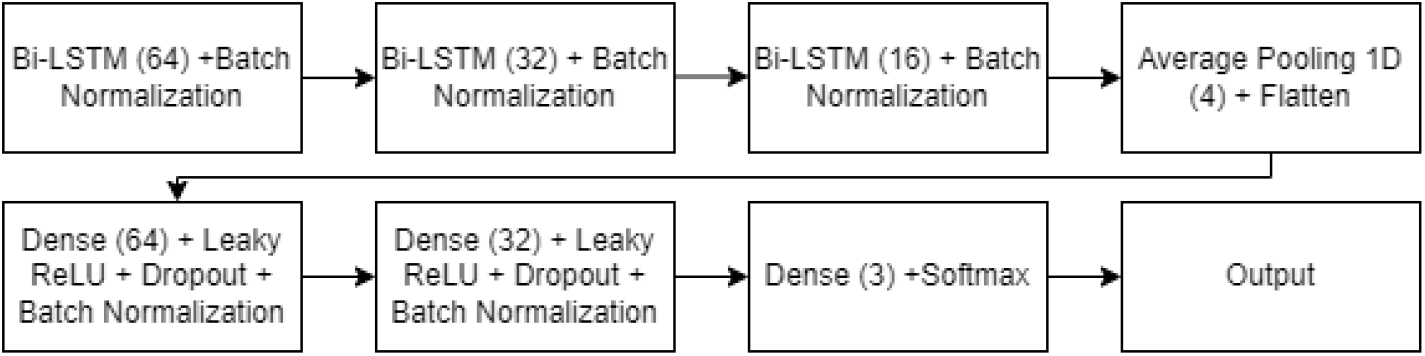
Diagram of the neural network architecture used for this study.

**Figure 2:**
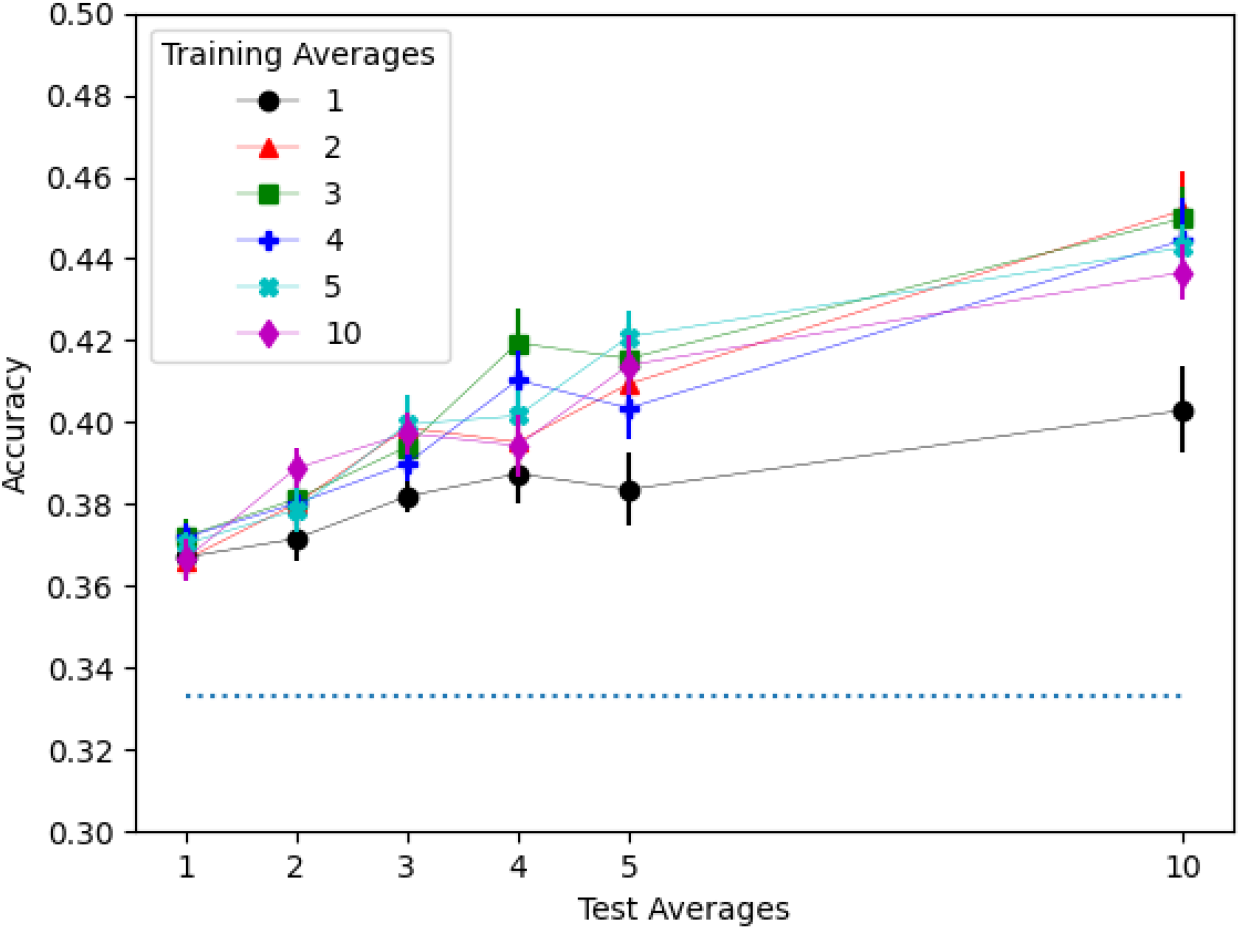
Accuracy by training average and test average. There were significant positive effects of increasing both the number of training and test averages. Error bars in all images reflect the standard error of the mean. Chance accuracy of 0.33 is represented by a dashed line in all images.

### 2.4. Training and Testing Paradigms

#### Training Averages Within Classes

We explored the effects of training the network on data averaged in varying amounts. In particular, we evaluated networks trained with 1, 2, 3, 4, 5, or 10 averaged samples and tested on data consisting of a single signal or 2, 3, 4, 5, or 10 averaged samples. The upper limit was set at 10 averaged samples due to data availability, while 6–9 averaged samples were omitted to limit the quadratic growth of the search space. When necessary to distinguish this training paradigm from other approaches described below, we will refer to it as the “simple” averaging paradigm.

We also explored the effects of intermixing various levels of averaging during the training of the network. In this training paradigm, each minibatch contained an equal proportion of training samples created through the averaging of 1, 2, 3, 4, or 5 raw samples. Henceforth, we will refer to this paradigm as the “intermixed” averaging paradigm.

#### Mixup and n-Mixup

We also explored the Mixup model for data augmentation. Mixup is a method for the weighted averaging of two signals and their labels, in which a beta distribution is sampled to determine the weighting of the average. Notably, the two signals are not required to be drawn from the same class. We chose an alpha parameter of 0.4, in line with the value used in the original paper [1]. We further explored an extension of Mixup that we refer to as n-Mixup, in which more than two signals can be combined by sampling from the Dirichlet distribution, which is the multidimensional generalization of the beta distribution. The parameters for the Dirichlet distribution are all set to 1, as in previous work [17], to produce a random sample from an n-1 dimensional simplex. As with the simple averaging approach described above, this training paradigm was also tested on data consisting of either a single signal or 2, 3, 4, 5, or 10 averaged signals.

#### Training and Testing Structure

A 60-20-20 train-validation-test split was employed with 20 iterations of random subsampling cross-validation per training average, with each level of test averaging being computed on the same split. We trained the model across subjects rather than within a single subject due to the limited availability of data per subject. Training averages were calculated at runtime through random selection. Similarly, when the test set included averages, those were randomly generated at test time, evenly distributed between the three classes. The test set of individual samples included approximately 263 trials per class, and a matching number of averaged test samples were generated when appropriate.

## 3. Results

In this section, classification results obtained from using 10 training averages and/or 10 test averages are included in the table and figures to demonstrate the continuation of the patterns taken to a more extreme degree, but left out of the statistical calculations due to the effect that the discontinuity between 5 and 10 averages would have on their interpretation.

Table 1 contains the average accuracies obtained over 20 iterations through the train-test process for all numbers of training and testing averages used in the simple averaging paradigm. All training and test configurations obtained above-chance accuracy. Training and testing on a single sample can be seen as the base case; this case achieved a 36.7% average accuracy. A one-way ANOVA over all numbers of training averages, considered at only one test average, showed no effect of training averages on accuracy (p *>* 0.7). Thus, it appears that simple averaging on training samples alone is insufficient to increase performance on relatively noisy single-trial test samples, at least in the current dataset.

**Table 1:** Training and Test Averages and Standard Deviations

| Training<br>Averages | Test Averages |  |  |  |  |  |
| --- | --- | --- | --- | --- | --- | --- |
|  | 1 | 2 | 3 | 4 | 5 | 10 |
| 1 | 36.7 (2.3) | 37.2 (2.3) | 38.2 (1.8) | 38.7 (3.2) | 38.4 (4.0) | 40.3 (4.7) |
| 2 | 36.6 (1.4) | 38.0 (2.3) | 39.9 (1.9) | 39.5 (3.2) | 40.9 (3.5) | 45.2 (4.2) |
| 3 | 37.2 (1.8) | 38.1 (3.3) | 39.4 (2.6) | 41.9 (3.8) | 41.6 (3.8) | 45.0 (3.5) |
| 4 | 37.2 (1.7) | 38.0 (2.5) | 39.0 (2.0) | 41.0 (3.2) | 40.3 (3.4) | 44.5 (4.5) |
| 5 | 37.0 (1.3) | 37.9 (2.5) | 39.9 (3.3) | 40.2 (2.7) | 42.1 (2.9) | 44.3 (2.6) |
| 10 | 36.6 (2.4) | 38.9 (2.2) | 39.7 (2.2) | 39.4 (3.5) | 41.4 (3.3) | 43.7 (3.0) |

However, a 5 (training averages: 1, 2, 3, 4, or 5) × 5 (test averages: 1, 2, 3, 4, or 5) two-way ANOVA over all combinations of numbers of training and test averages showed a significant effect of training averages (F(4, 475) = 6.401, p < 0.001) and test averages (F(4, 475) = 32.493, p < 0.001), with no interaction between those factors (p *>* 0.2). Furthermore, there were significant linear (p < 0.001) and quadratic (p = 0.005) trends for the training averages factor. Both of these trends were largely driven by the difference between training on a single trial versus any number of averaged trials. There was also a significant linear trend for the number of test averages (p < 0.001), with more averages yielding higher accuracy. Figure 3 demonstrates the effect of any amount of averaging training data (by combining the results from 2 through 5 training averages) compared to training on a single trial, at all numbers of test averages. As can clearly be seen in the figure, the benefit of more test averaging was greater when also employing averaging during training.

**Figure 3:**
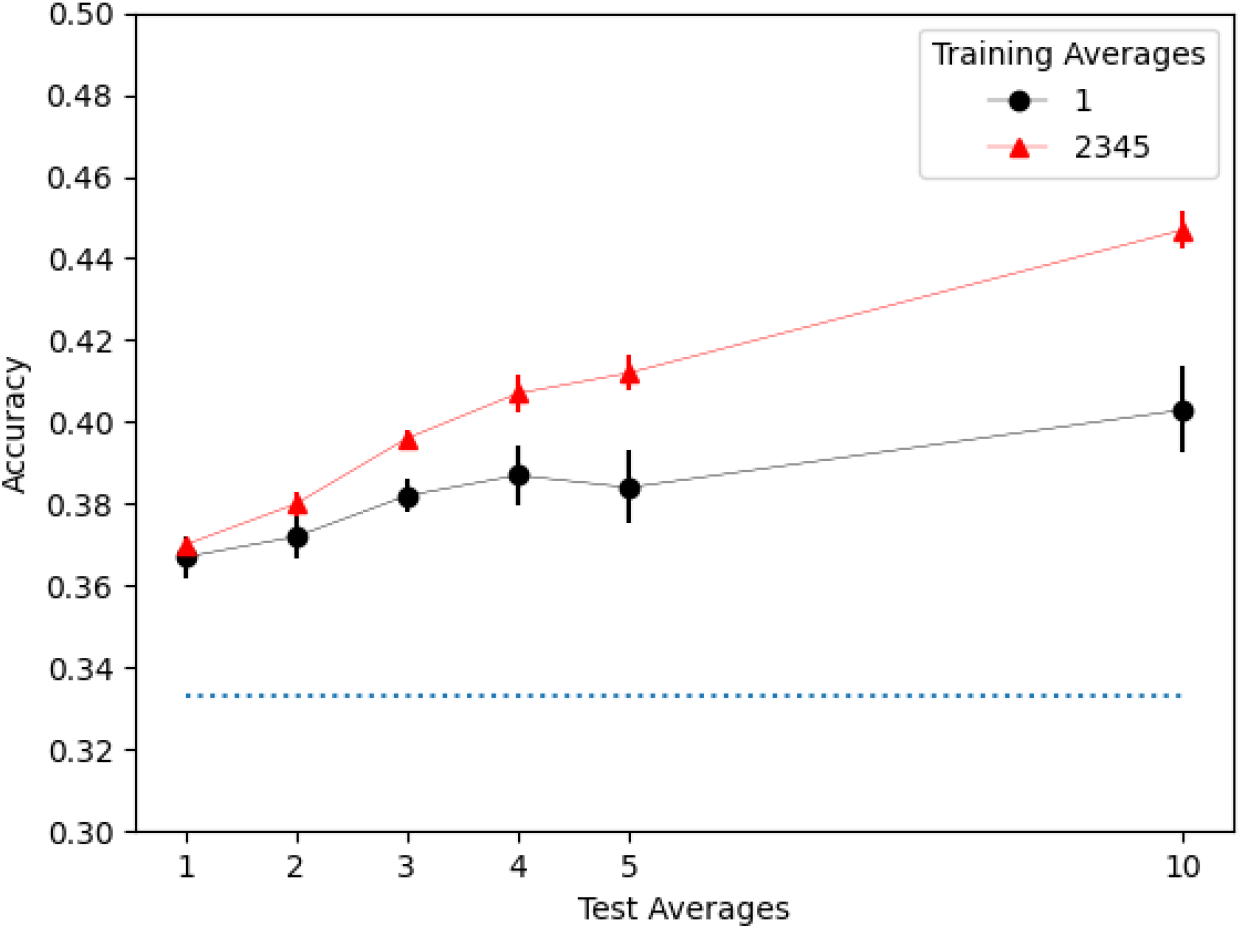
One training average performed significantly worse than any other number of training averages, though both benefited from increasing the number of test averages.

We also performed a 2 (training paradigm: simple averaging versus intermixed) × 5 (test averages: 1, 2, 3, 4, or 5) two-way ANOVA to assess differences between those training paradigms. As in the previous analysis and Figure 3, the simple averaging condition was calculated by combining the results from 2 through 5 training averages. Intermixing did not significantly differ from simple averaging as a main effect of training paradigm (p *>* 0.7). There was still a significant main effect of test averages (F(4, 190) = 13.815, p < 0.001), but no interaction between the two factors, p *>* 0.2. See Figure 4.

**Figure 4:**
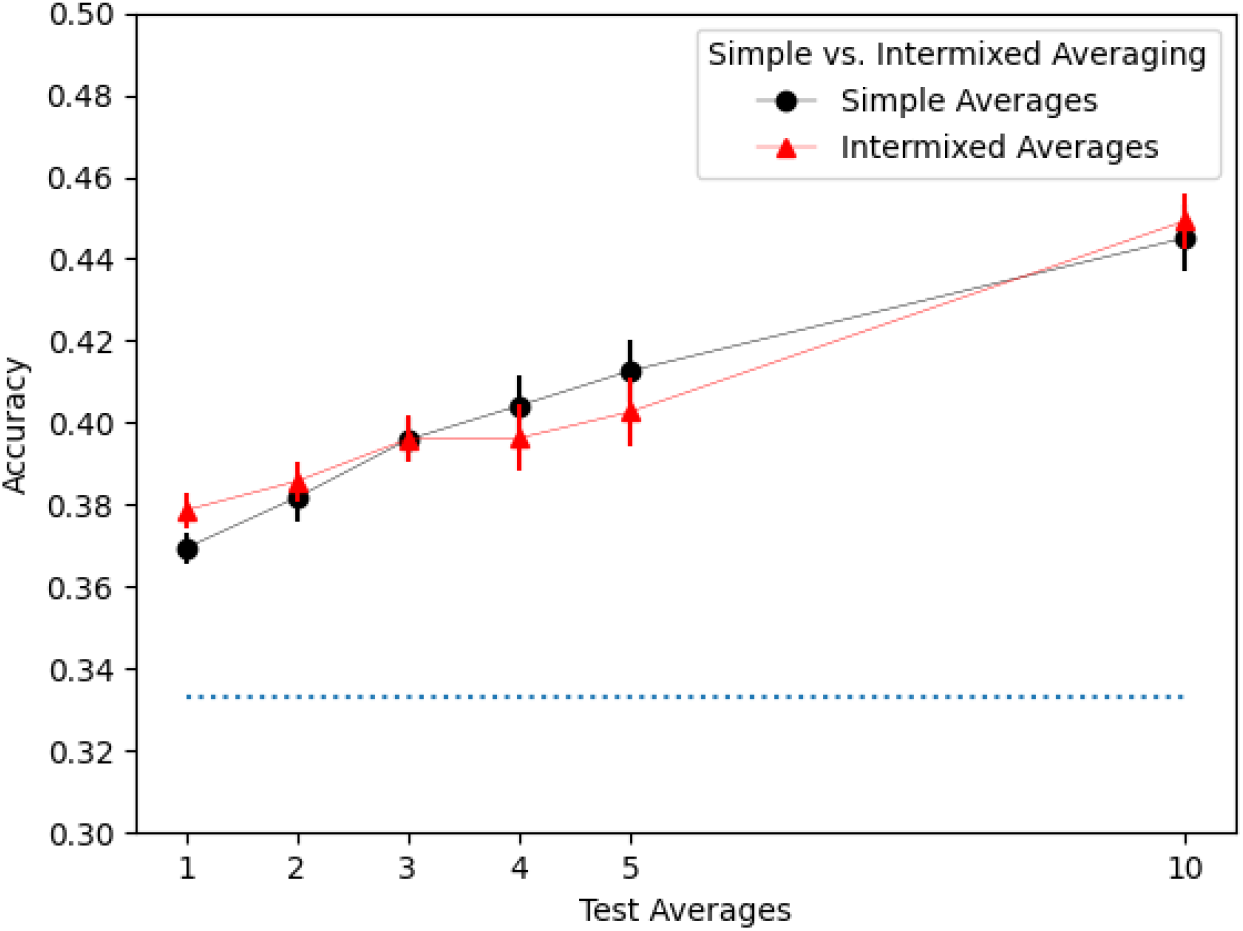
Training with intermixed averages per sample did not show a significant difference from training with a static number of averages per sample.

We next performed a 4 (training averages: 2, 3, 4, or 5) × 5 (test averages: 1, 2, 3, 4, or 5) two-way ANOVA on the Mixup and n-Mixup training paradigms. This demonstrated a significant main effect of training averages (i.e., the number of samples used for performing Mixup, in this case) for standard (2-average) Mixup and n-Mixup at 3, 4, and 5 averages (F(3,380) = 8.861, p < 0.001). This pattern was characterized by higher accuracy in standard Mixup and 3-Mixup versus lower performance in 4- and 5-Mixup. In other words, increasing the number of training averages did not benefit Mixup and in fact was detrimental to performance. There was a significant main effect of test averages (F(4, 380) = 3.781, p = 0.005), with somewhat better performance at higher numbers of test averages. The interaction between testing and training averages was not significant (p *>* 0.6). See Figure 5.

**Figure 5:**
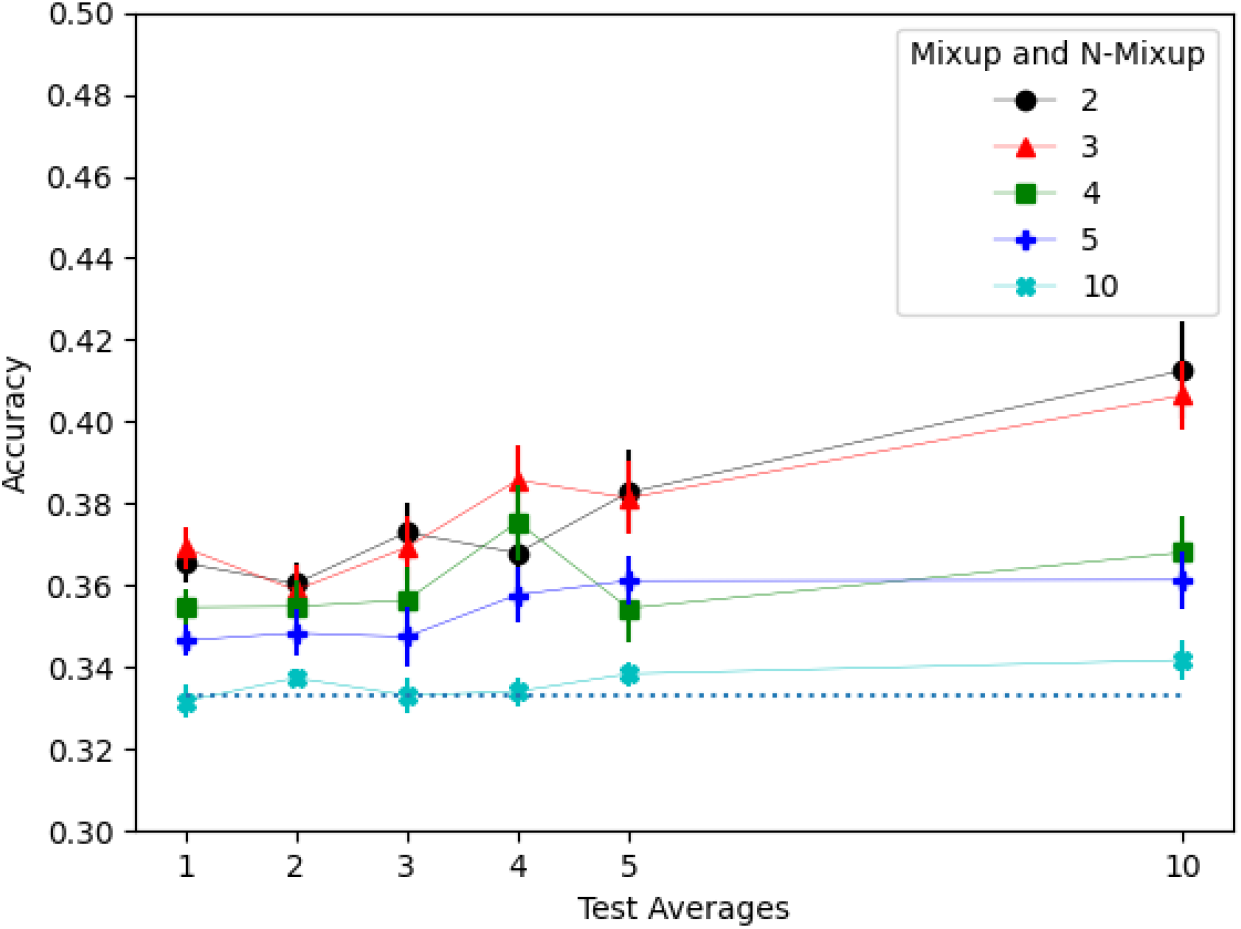
Mixup generally performed poorly, with a greater degradation when using more training averages.

We then compared simple averaging and n-Mixup with a 2 (training paradigm) × 4 (training averages: 2, 3, 4, or 5) × 5 (test averages: 1, 2, 3, 4, or 5) three-way ANOVA. This analysis showed a highly significant main effect of training paradigm (F(1, 760) = 192.313, p < 0.001), with simple averaging techniques performing better. There were also main effects of both training averages (F(3, 760) = 5.865, p < 0.001) and test averages (F(4, 760) = 25.897, p < 0.001). There was additionally an interaction between training paradigm and training averages (F(3, 760) = 5.282, p = 0.001), largely reflecting the poor performance of 4- and 5-Mixup, as well as an interaction between training paradigm and test averages (F(4, 760) = 6.439, p < 0.001), reflecting the fact that n-Mixup benefited less from increasing the number of test averages than simple averaging did. The interaction of training and test was not statistically significant (p = 0.13), nor was the interaction of training, type, and test (p*>*.99). Figure 6 shows standard Mixup plotted against the simple averaging technique; for easier viewing, the simple averaging condition was once again calculated by combining the results from 2 through 5 training averages in this figure.

**Figure 6:**
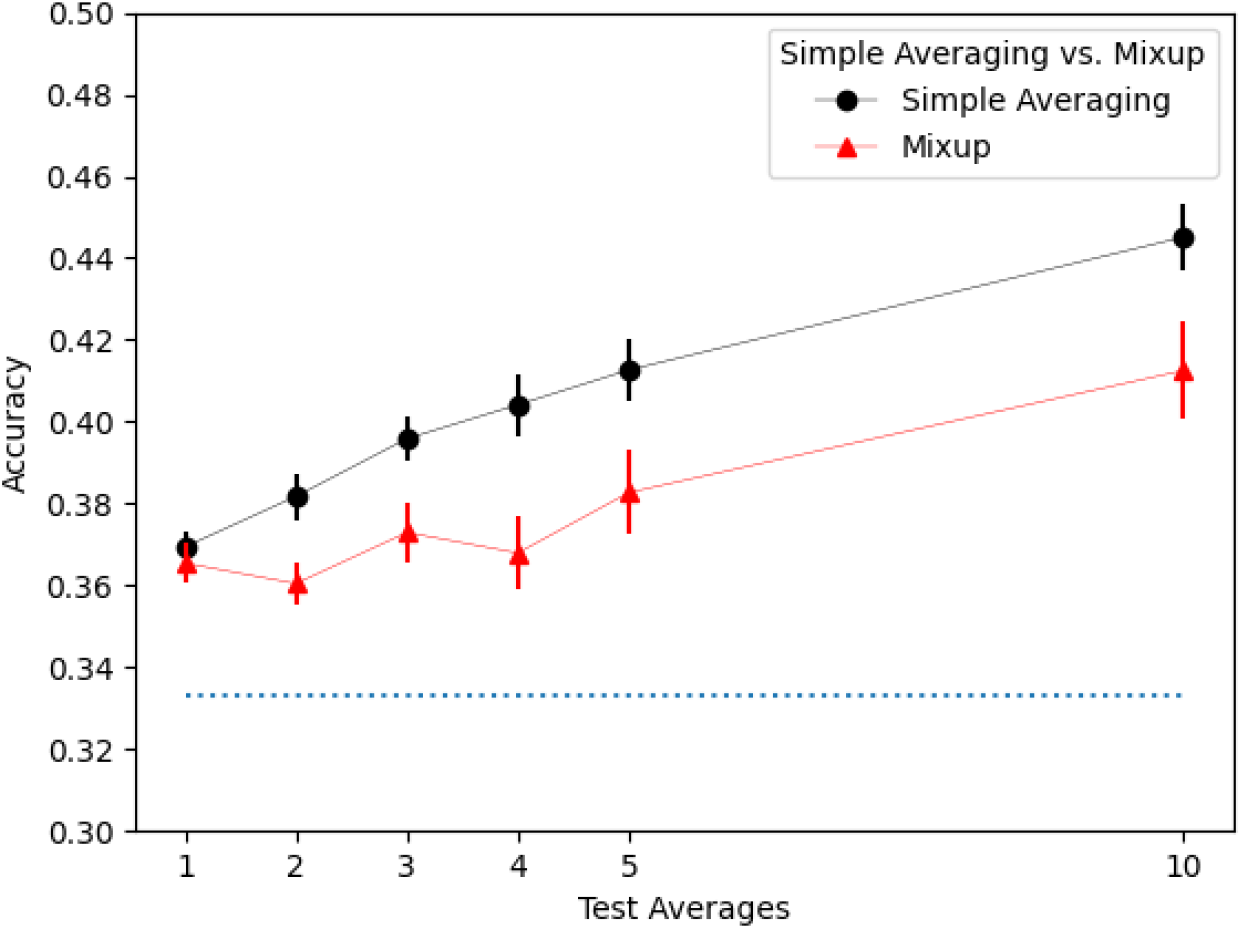
Standard Mixup performed worse than simple averaging.

To examine differential effects of averaging on the classifiability of different stimulus classes, we then calculated per-class F1 scores for the simple averaging training paradigm separately for each category. Figures 7, 8, and 9 show the F1 score of each class (recalled faces, scenes, and words respectively) at each level of training and test averaging. We analyzed each of those sets of F1 scores with its own 5 (training averages: 1, 2, 3, 4, or 5) × 5 (test averages: 1, 2, 3, 4, or 5) two-way ANOVA.

**Figure 7:**
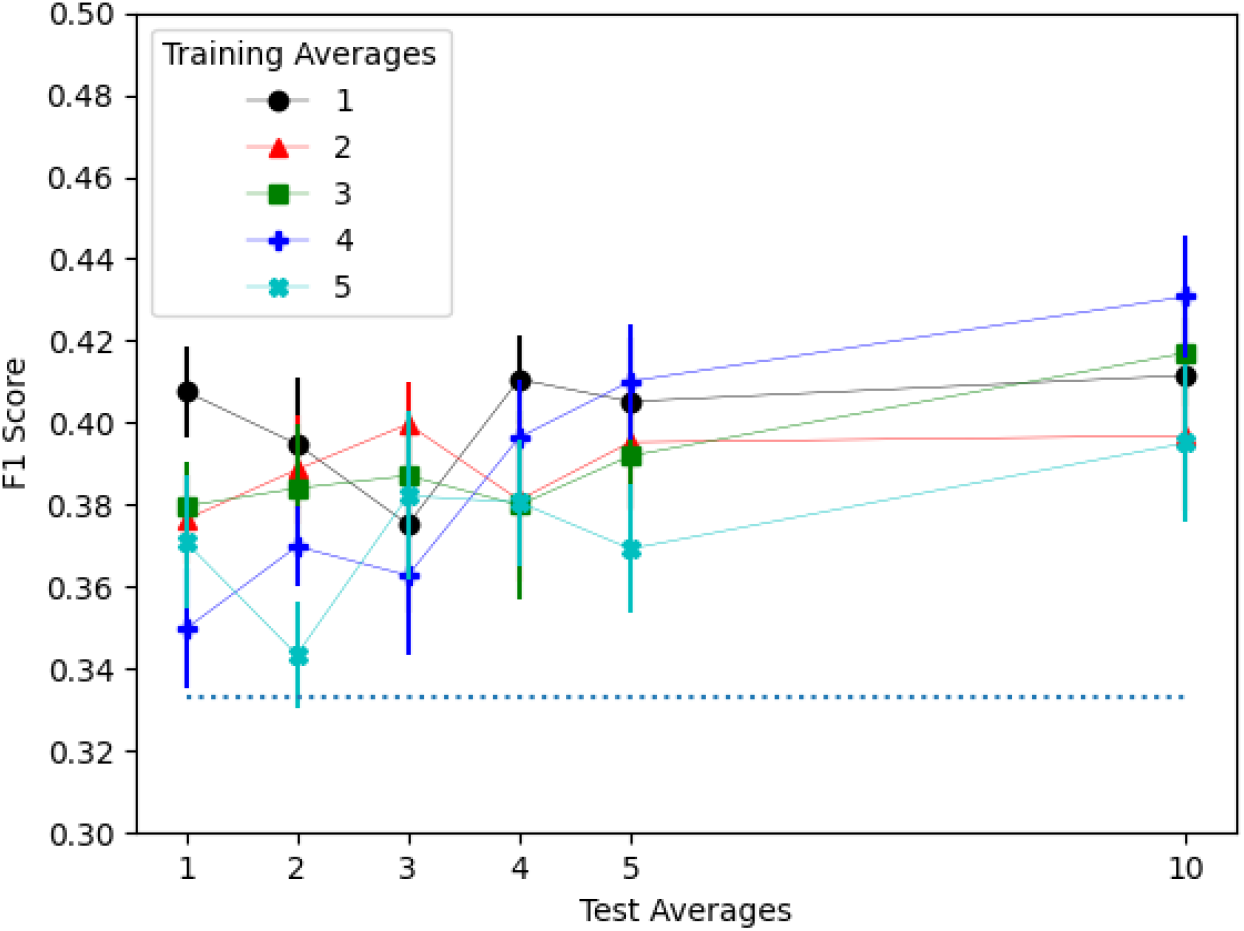
Face F1 scores. Unlike other categories of images, increasing the number of training averages decreased the performance on classifying recalled faces. There was no significant effect of test averages.

**Figure 8:**
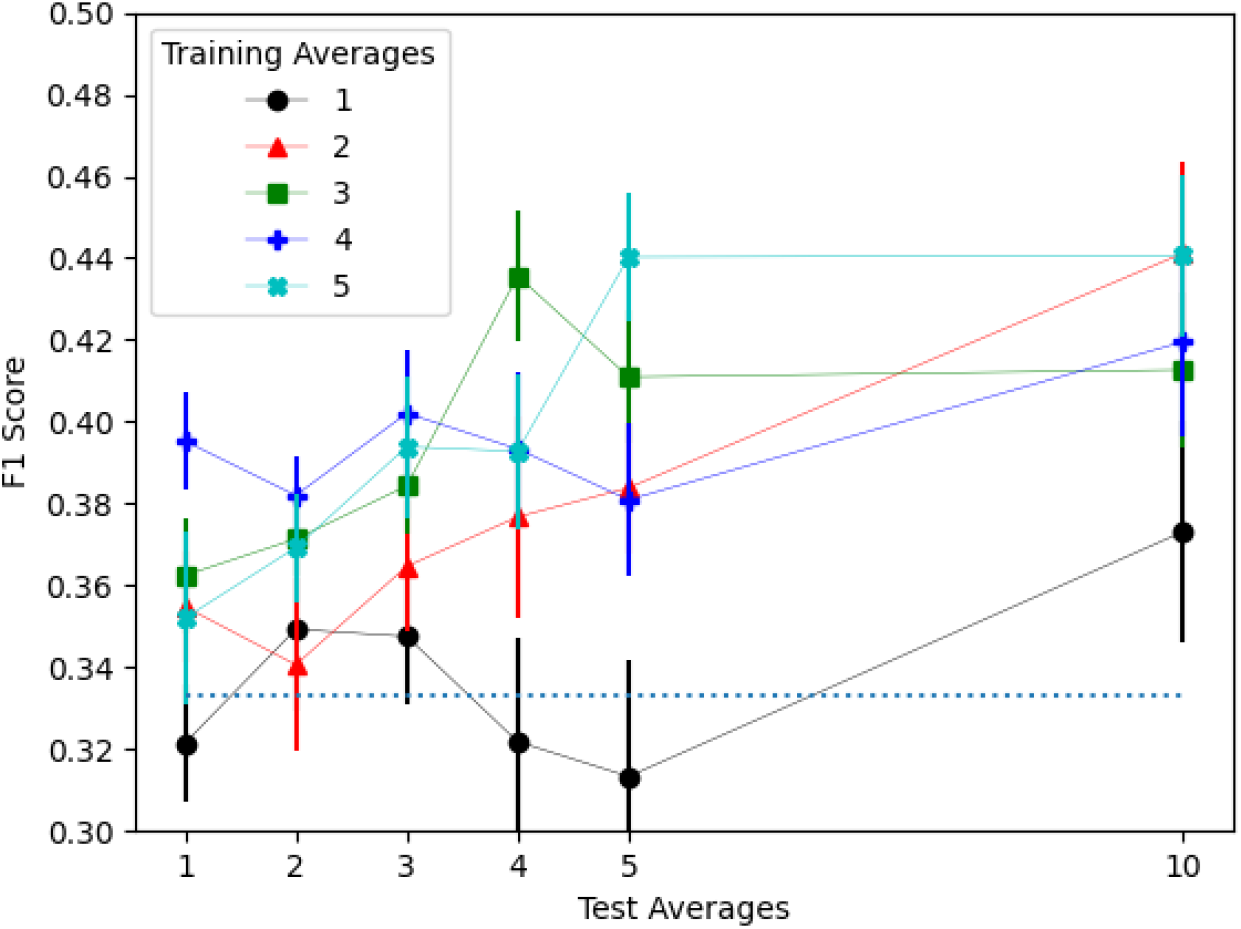
Scene F1 scores. Increasing the number of both training and test averages increased performance in classifying recalled scenes.

**Figure 9:**
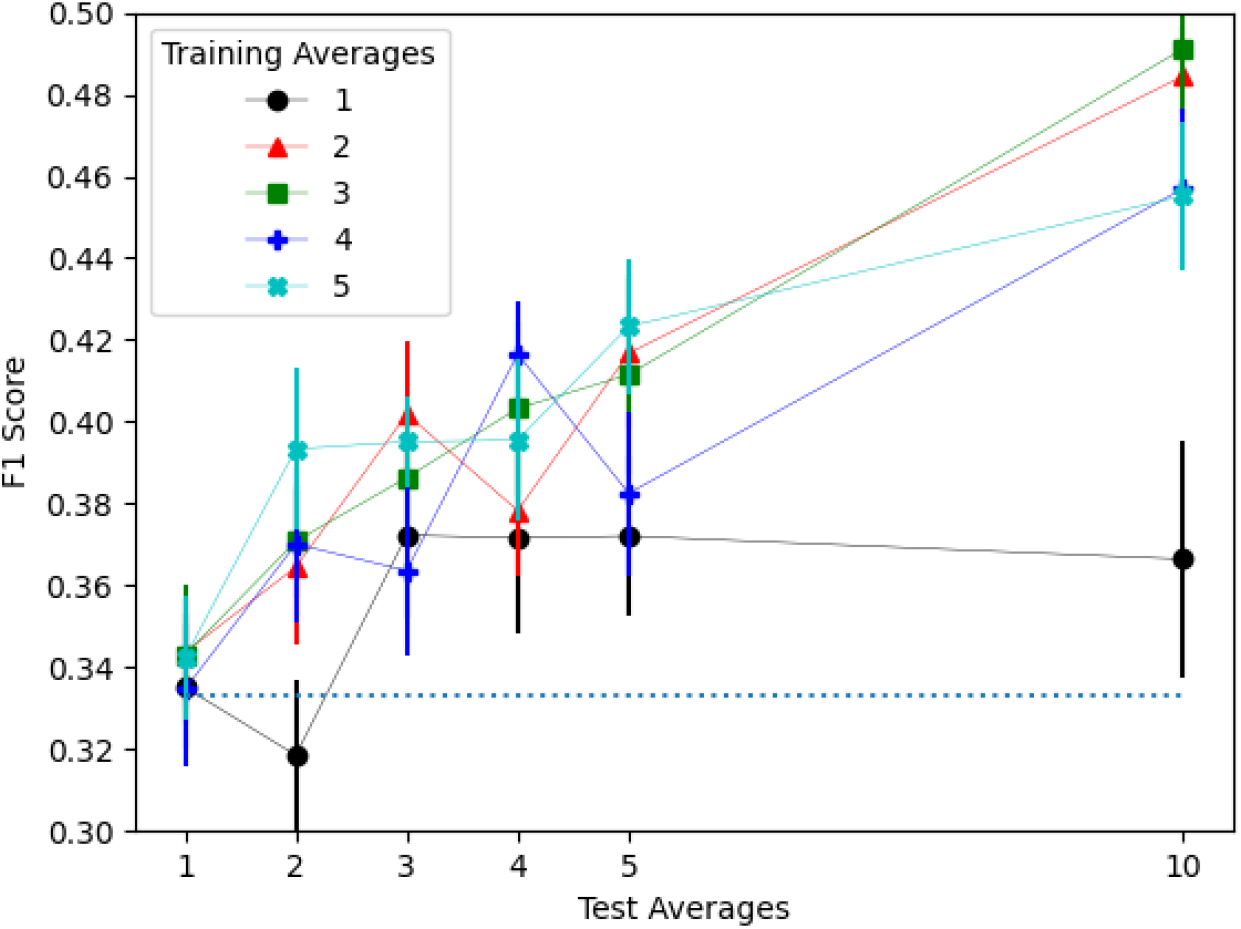
Word F1 scores. Increasing the number of both training and test averages increased performance in classifying recalled words.

Recalled face images showed a significant main effect of training averages (F(4, 475) = 2.549, p = 0.039), but no main effect of test averages (p *>* 0.2) or interaction of the two (p *>* 0.5). There was a significant linear trend for training averages (p = 0.002), with higher training averages leading to worse classification results, which notably was the opposite of the overall pattern we observed with all of the classes combined.

Recalled scenes showed a significant main effect of both training averages (F(4, 475) = 11.541, p < 0.001) and test averages (F(4, 475) = 2.731, p = 0.029), but no interaction (p *>* 0.1). There was a significant linear trend for training averages (p < 0.001) as well as a significant quadratic trend (p < 0.001). These trends appeared to be driven by the fact that more training averages produced higher F1 scores, but this effect reached an asymptote around 3 training averages. Additionally, there was a significant linear trend for test averages (p = 0.002) with more test averages monotonically increasing performance.

Recalled words showed a significant main effect of both training averages (F(4, 475) = 2.909, p = 0.021) and test averages (F(4, 475) = 9.418, p < 0.001), but no interaction (p *>* 0.7). There were significant linear trends for both training averages (p = 0.012) and test averages (p < 0.001), with more averages producing better performance in both cases.

## 4. Discussion

The most clear result we observed in this study was the positive impact of averaging the data at test time. In most of the paradigms explored, there was a significant positive linear effect of increasing the number of samples averaged during testing, up to the maximum value of 10 tested in this dataset. One notable exception was within the recalled face class, which did not experience a positive effect of trial averaging at test time, though there did not appear to be a negative effect of averaging at test time either. Thus, averaging test data is likely advisable when compatible with the aims and experimental design of a particular study.

There was an overall positive effect of averaging within a class during training, though the benefit of increasing amounts of averaging was not as apparent or consistent as it was for averaging during test. Again, there was a notable exception in that, within recalled faces, more averaging actually resulted in a decrease in F1 score.

The differences seen in the effects of averaging between recalled faces and recalled scenes or words is potentially due to the universal nature of the models trained. That is, there may be higher person-to-person variance in how faces are remembered than how scenes or words are remembered, leading to averaging samples between subjects having a destructive effect on the overall signal. It is well understood, for example, that people are able to remember the faces of other people sharing their race better than those of differing races [18]. Future studies may also wish to consider these issues on a per-stimulus-category basis, but in the present study, the negative effects of training averaging on faces were still not enough to offset the overall benefit; thus, we would expect future studies to experience an overall improvement in classification performance from using averaging during training even if not controlling for effects of stimulus category.

Another interesting result was that training with a fixed amount of averaging (e.g., averaging exactly two or exactly three samples for the entire training process) is at least as effective as intermixing the number of samples averaged for each datum included in a mini-batch. *A priori*, we had conjectured that learning on both the noisier individual samples and higher-signal averaged samples would have allowed for the best of both worlds in being both robust to noise and having clearer signals to train on, but this turned out not to be the case.

Also somewhat contrary to our initial expectations, all forms of Mixup performed worse than simple, within-class averaging. It seems less likely that this was due to the differences in the weighting of these paradigms (equal weighting per sample for simple averaging versus potentially unequal for Mixup) and more likely that it was due to the difference between within-class and between-class averaging. This suggests that EEG signal classification may not experience the same benefit of superimposition that has previously been found in studies of classifying optical images. That is, averaging samples and their labels from different classes may have a destructive effect on the overall signal, reducing the ability of the model to learn. Further support for this idea comes from the fact that higher n’s of n-Mixup showed increasingly worse performance.

There are a few clear avenues for future research. Most immediately, the effects studied here only represent a single dataset. Expanding the work to include more datasets would costly given the sheer number of models trained, but will be important for stronger claims of generalizability. With the differing pattern of the effects of averaging seen in the classification of recalled face EEG signals versus recalled scenes or words, further studies of when averaging is appropriate for certain categories of stimuli or certain experimental designs would be beneficial. Another path for future research would be exploring larger datasets with enough data per subject to train individualsubject models. Again, with the effects we observed in the classification of recalled face EEG signals, alongside the large individual differences that can be seen in studies of face processing and short-term memory, it seems likely that the patterns of effects in a within-subject model would be different from those we observed in the between-subjects models used in the present study.

## Acknowledgements

This work was supported by NSF/EPSCoR Grant 1632849, RII Track-2 FEC: Neural networks underlying the integration of knowledge and perception, awarded to MJ and colleagues.

## Notes

### Competing Interest Statement

The authors have declared no competing interest.

